# Environmental correlates and functional consequences of bill divergence in island song sparrows

**DOI:** 10.1101/2021.09.15.460486

**Authors:** Maybellene P. Gamboa, Cameron K. Ghalambor, T. Scott Sillett, W. Chris Funk, Ross A. Furbush, Jerry F. Husak, Raymond M. Danner

## Abstract

Inferring the environmental selection pressures responsible for phenotypic variation is a challenge in adaptation studies as traits often have multiple functions and are shaped by complex selection regimes. We provide experimental evidence that morphology of the multifunctional avian bill is related to climate, not foraging efficiency, in song sparrows (*Melospiza melodia*) on the California Channel Islands. Our research builds on a study in song sparrow museum specimens that demonstrated a positive correlation between bill surface area and maximum temperature, suggesting a greater demand for dry heat dissipation in hotter, xeric environments. We sampled contemporary sparrow populations across three climatically distinct islands to test the alternate hypotheses that song sparrow bill morphology is either a product of vegetative differences with functional consequences for foraging efficiency or related to maximum temperature and, consequently, important for thermoregulation. Measurements of >500 live individuals indicated a significant, positive relationship between maximum temperature and bill surface area when correcting for body size. In contrast, maximum bite force, seed extraction time, and vegetation on breeding territories (a proxy for food resources) were not significantly associated with bill dimensions. While we cannot exclude the influence of foraging ability and diet on bill morphology, our results are consistent with the hypothesis that variation in song sparrows’ need for thermoregulatory capacity across the northern Channel Islands selects for divergence in bill surface area.

**SUMMARY STATEMENT:** Island song sparrow bill differences are correlated with climate, not vegetation, and experimental evidence finds no functional effect on foraging efficiency. This suggests many factors shape this multifunctional trait.

## INTRODUCTION

Determining the environmental factors that drive adaptation in traits is a central goal in evolutionary biology, but this is often challenging in natural populations (Kawecki and Ebert, 2004; MacColl, 2011; Reznick and Travis, 1996). Such challenges arise because traits may serve different functions such that the observed phenotypic variation is a product of multifarious selection pressures (e.g., Egea-Serrano et al., 2014; Pfrender, 2012; Shultz and Burns, 2017; Templeton and Shriner, 2004; Wilkins et al., 2013). Multiple selection pressures can act either synergistically, shifting the population phenotypic mean towards a predictable adaptive optimum, or act antagonistically such that the observed phenotypic means represents a compromise, or trade-off between different functions (Svensson and Calsbeek, 2012). This concept of adaptation as a compromise between different functions is reinforced by empirical studies of natural populations (e.g., Egea-Serrano et al., 2014; Ghalambor et al., 2003; Kim et al., 2011; Robinson et al., 2006). Thus, testing which aspects of the environment act as important selection pressures requires consideration of the different functions of a given trait and the functional consequences associated with shifting trait means (Ghalambor et al., 2003; Jones et al., 1977).

The avian bill is one of the most studied multifunctional, morphological traits. The bill is involved in many fitness-related behaviors including ectoparasite removal (Clayton et al., 2005), communication (Ballentine, 2006; Podos, 2001), tool creation and use (Fayet et al., 2020; Rutz et al., 2016; Troscianko et al., 2012), thermoregulation (Greenberg et al., 2012; Ryeland et al., 2017; Symonds and Tattersall, 2010), and, most notably, food acquisition (Barbosa and Moreno, 1999; Benkman, 1993; Temeles and Kress, 2003). Consequently, predicting local optima for bill sizes is difficult given the potentially conflicting functional demands. For example, an increase in bill morphology in the Darwin’s finches is associated with improved foraging efficiency on hard seeds, yet it is also predicted to cause correlated changes in syllable rate and frequency bandwidth of vocal signals, which alters song production (Podos and Nowicki, 2004). Furthermore, finches with increased bill surface area have greater heat dissipation, which is hypothesized to improve thermoregulatory function (Tattersall et al., 2018). Similar interspecific patterns of bill divergence correlated with multiple environmental drivers and resulting in functional consequences have been documented in other passerines as well (Friedman et al., 2019). Bill morphology in any bird species is, therefore, a product of trade-offs among multiple selection pressures including, but not limited to, climate, food resources, and vocal signaling. Bill dimensions also have a strong genetic component, indicating that this important trait can readily evolve in response to selection (Åkesson et al., 2008; Boag, 1983; Grant, 1983; Jensen et al., 2003; Keller et al., 2001). Given that the strength of selection may shift over time and space (Siepielski et al., 2009; Siepielski et al., 2013), investigating avian bill morphology differences among environments and populations can provide insight into how multiple selection pressures act to generate and maintain variation.

The relationship between bill morphology and foraging ability has received extensive attention, with numerous empirical studies finding correlations between bill size and characteristics of available food resources or foraging ability (e.g., Langin et al., 2015; Nebel et al., 2005; Temeles et al., 1993). For instance, bill depth in the medium ground finch (*Geospiza fortis*) is positively correlated with the abundance of large, hard seeds, and evolution in response to fluctuations in seed availability across years results in rapid adaptation (Grant and Grant 2006). Relatively small modifications in bill morphology among Darwin’s finches result in functional differences in bite force (Herrel et al., 2010). This strong selection pressure on bill morphology for improved foraging ability has resulted in diversification and adaptive radiation in several avian families (Benkman, 2003; Burns et al., 2003; Grant and Grant, 2002; Lamichhaney et al., 2015; Lerner et al., 2011; Parchman et al., 2006). These striking results coupled with other empirical studies suggest bill morphology should be strongly associated with foraging and dietary resources. Yet, selection for foraging efficiency may not operate in isolation from other environmental and ecological drivers.

The avian bill has also been studied in the context of thermoregulation and, specifically, heat dissipation (Tattersall et al., 2016). The bird bill is an exposed, vascularized network that exchanges heat directly with the environment, thereby acting as a ‘thermal window’ between internal temperature and external, ambient temperature (Hagan and Heath, 1980; Symonds and Tattersall, 2010; Tattersall et al., 2009). Increased blood flow to the vascularized region of the bill results in increased heat dissipation (Tattersall et al., 2016). By dissipating dry heat through radiation rather than panting, birds in arid, xeric environments may reduce evaporative water loss while maintaining body temperature equilibrium (Dawson, 1981; Tattersall et al., 2016). However, selection for large bills to increase thermoregulatory capacity could also impact diet depending on the availability of food resources and on how strongly bill dimensions affect functionality, namely in bite force and seed extraction (Herrel et al., 2010; Soons et al., 2015; van der Meij and Bout, 2004). Thus, testing the relative importance of food resources and climate on bill variation and evaluating the functional consequences of population shifts in bill morphology allows for inferring how selection operates on integrated traits.

Here, we investigate the relationship between variation in bill surface area, feeding performance, and climate in song sparrows (*Melospiza melodia*) breeding on the California Channel Islands. These island populations provide a model system for investigating how environmental factors influence adaptation in bill morphology. Song sparrows are continuously distributed along a climatic gradient ranging from cold, wet, and very windy on San Miguel and Santa Rosa Islands to hot, arid, and less windy on Santa Cruz and Anacapa Islands (Schoennerr et al., 1999; Fig. 1). In this system, maximum island temperature has been shown to be positively correlated with bill surface area of song sparrow museum specimens and implicated in aiding in thermoregulation (Greenberg and Danner, 2012). Yet, the islands’ west-to-east climate gradient is also associated with vegetation differences (Junak et al., 2007) that could affect song sparrow habitat composition and food availability. We sampled contemporary sparrow populations across three climatically distinct islands (San Miguel, Santa Rosa, and Santa Cruz Islands) to test two hypothesized functional adaptations in song sparrow bills: thermoregulation and foraging efficiency. If bill morphology is acting as a thermoregulatory trait, we predicted that variation among song sparrow populations would be correlated with climatic differences across islands. If foraging efficiency explained variation in bill morphology, we predicted that bite force or seed extraction time would change as a function of bill dimensions, given that song sparrows primarily consume seeds during the non-breeding season (Arcese et al., 2002). We additionally assessed plant composition in song sparrow habitats (a proxy for food resources) across the three islands. Our combined assessments of the environmental correlates and functional consequences of bill variation allowed us to infer how complex selection regimes shape individual morphology and feeding performance.

**Figure 1.**
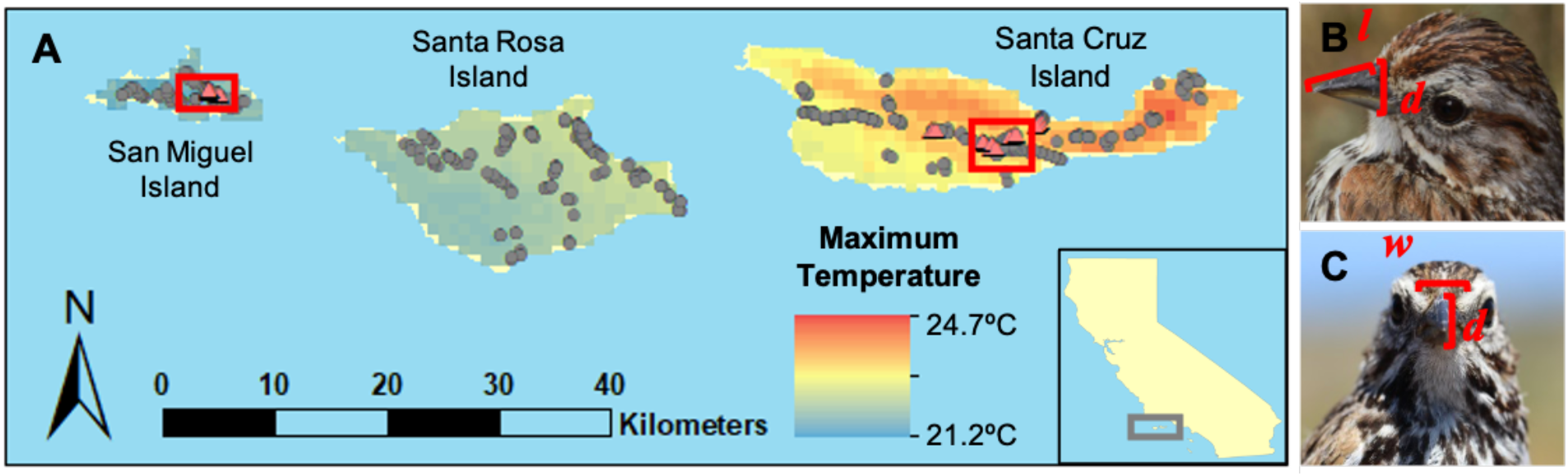
Sampling locations for comparison of song sparrow bill morphology (grey circles), seed extraction time (red triangles), and maximum bite force (red boxed regions) across three climatically-distinct islands (A) and measurements of bill length, depth (B), and width (C) used for quantifying bill morphology. Bill length (*l*), depth (*d*), and width (*w*) were taken from the anterior edge of the nares and used to calculate residual (body-size corrected) bill surface area following Greenberg and Danner (2012). All sampling was conducted during the breeding season (February-June) from 2014-2016. Inset shows the location of the northern Channel Islands with respect to California.

## MATERIALS & METHODS

### Animal capture and morphological measurements

We captured and measured song sparrows on San Miguel, Santa Rosa, and Santa Cruz Islands during three breeding seasons (February-June 2014-2016). All birds were target-captured using mist-nets and song playback in the morning (thirty minutes before sunrise to four hours after sunrise) when territory defense and foraging activities are high. We measured bill dimensions (depth, width, and length) from the anterior edge of the nares (Fig. 1B,C), tarsometatarsus length, wing length (0.01 mm precision), and mass. Estimates of bill depth, width, and length were used to generate total bill surface area, following Greenberg and Danner (2012). When comparing bill surface area among populations, we used the residual of a linear regression model with bill surface area as the response (bill_sc_) and the first principal component of an analysis of tarsometatarsus length and wing length (PC1_bod_) as the predictor (Greenberg and Danner, 2012). Tarsometatarsus and wing lengths are indicators of structural body size in birds, and this approach allows us to control for allometry (Rising and Somers, 1989). Negative values for residual bill surface area are indicative of smaller bill sizes than predicted by body sizes alone and, conversely, positive values suggest larger bill sizes than predicted by body sizes alone.

We applied nonparametric tests in R (R Core Team 2020) to determine whether raw (uncorrected for body size; bill_raw_) and residual (corrected for body size; bill_sc_) bill surface area differed by island. Specifically, we used Kruskal-Wallis tests (*kruskal.test*) to assess whether island raw and residual bill surface area mean ranks differed, while accounting for unequal variances. We did not include sex in our models, because body size corrections account for allometric differences between males and females. To further investigate island differences, we performed post-hoc, pairwise comparisons of island mean ranks and output 95% confidence intervals around estimated differences using Mann-Whitney U tests (*wilcox.test*) with Bonferroni corrections for multiple-testing.

### Testing if song sparrows experience different thermal environments and habitats

We extracted climate data at 1 km^2^ (30s) spatial resolution from WorldClim v. 1.4 (Hijmans et al., 2005) for all sampling locations using ArcGIS v. 10.4 (ESRI 2011) to test if birds on different islands experience different temperatures. Minimum, maximum, and mean monthly temperatures were highly correlated (*r^2^* > 0.7). Based on previous work showing a significant, positive correlation between residual bill surface area and maximum temperature (Greenberg and Danner, 2012), we limited our analyses to maximum temperature and extracted monthly temperatures in July, the hottest month of the year on average for the northern Channel Islands. Extracted maximum temperature values (*T*) served as a proxy for climate in individual song sparrow territories. As with analyses of bill dimensions, we performed a nonparametric Kruskal-Wallis test to compare island mean ranks and quantified pairwise differences in island mean ranks and 95% confidence intervals using post-hoc Mann-Whitney U tests.

Song sparrows generally occupy low shrubland and, occasionally, riparian and coastal sage scrub habitat across the Channel Islands (Shuford and Gardali, 2008). To infer if sparrows use different habitats with different plant species (a proxy for dietary resources), we conducted vegetation surveys within a 25-meter radius of each mist-net location for sampled birds. Because sampling occurred during the breeding season, these measurements were taken within the approximated territories of sampled birds and, thus, reflect the plant species available. We recorded dominant woody vegetation type to the species level, when possible, for all plants that comprised >50% of the total area. Additionally, we identified commonly occurring vegetation types and categorized the relative abundance of these vegetation types at all sparrow sampling sites. Presence and coverage of vegetation types within the sampling area was recorded using a ranked scale including absent (0%; 1), trace (<10%; 2), some (10-25%; 3), prominent (25-50%; 4), and dominant (>50%; 5). To infer island-level vegetative differences among song sparrow territories, we used Fisher’s exact test in R (*fisher.test*) to test for an association between ranked abundance of vegetation types and island. We modeled the null distribution of the test statistic using 10,000 Monte Carlo simulations, allowing us to estimate the p-value under the null hypothesis that the abundance of different vegetation types is independent of the island sampled.

We modeled vegetation for individual sampling locations by transforming ordinal vegetation data to quantitative dimensions using nonlinear principal components analysis (NLPCA) implemented in the R package *Gifi* (Mair and de Leeuw, 2019). This multivariate method reduced the complexity of correlated vegetation variables to two principal components in ordination space while accounting for ranked abundance of each vegetation type. We visually inspected NLPCA results and assessed loadings of categorical values on dimensional space to infer what factors drive variation in the first two axes of variation (PC1_veg_ and PC2_veg_) among song sparrow territories. We plotted individual sampling locations in vegetation space along PC1_veg_ and PC2_veg_ axes and constructed 95% kernel density contours to visually assess overlap among islands in vegetation space. To statistically compare these reduced vegetation descriptors (PC1_veg_ and PC2_veg_) among islands, we again applied a nonparametric Kruskal-Wallis test to account for unequal variances. We performed post-hoc Mann-Whitney U tests with Bonferroni corrections and extracted estimated island means and 95% confidence intervals around these differences. NLPCA dimensions (PC1_veg_ and PC2_veg_) were used for subsequent tests relating habitat and residual bill surface area.

We performed linear regression to determine whether vegetation and climate were significant predictors of residual bill surface area. We used the R function *lm* to model residual bill surface area as predicted by maximum temperature and two vegetation dimensions (PC1_veg_ and PC2 _veg_) resulting from NLPCA of all vegetation sampling sites. We generated 95% confidence intervals around unstandardized beta estimates using the function *confint* in the base *stats* package in R and extracted standardized beta coefficients for all predictors using the R package *lm.beta* (Behrendt, 2014). Unstandardized and standardized beta coefficients allowed us to evaluate the relative importance of climate versus vegetation (a proxy for diet) for predicting variation in song sparrow bill surface area.

### Measuring maximum bite force to infer the functional consequences of bill variation

To determine whether divergence in bill morphology results in functional consequences for food acquisition, we compared bill functional morphology between Santa Cruz and San Miguel Island song sparrows. We expected functional differences to be largest between these populations based on pronounced climate differences between islands and on phenotypes observed in museum specimens by Greenberg and Danner (2012). All sampling and estimates of bite force were performed in early spring 2014, when birds are primarily foraging on seeds. We measured maximum bite force in the field using a custom-manufactured force meter (Herrel et al., 2005; van der Meij and Bout, 2004). Briefly, we used a piezoelectric isometric force transducer (type 9203, Kistler, Switzerland) fitted to custom-built stainless-steel bite plates (specifications in Herrel et al., 1999) and connected to a charge amplifier (type 5995, Kistler, Switzerland). A micrometer head allowed adjustment of the spacing between bite plates. For each measurement, we held the bird upright and positioned the bite force meter between the mandible and maxilla. We positioned the ends of the plates two-thirds of the distance from the bill tip to commissure, the location where song sparrows crush seeds (Danner, *pers. obs*.). The meter had a precision of 0.1 Newtons. We recorded the maximum bite force over a period of 15 seconds. Birds were gently coaxed to open the bill by tapping on the tomia with a thin, metal spade. Preliminary analyses on song sparrows indicated that bite force did not decline across observations when measurements were interspersed with 15-second rest intervals (Danner, *pers. obs*). Thus, we recorded three measurements per bird with one-minute rest intervals to ensure recovery and used the maximum bite force for all further analyses. All morphological measurements were taken shortly before releasing the birds to minimize the effect of handling stress on bite force and seed extraction trials. We maintained the same force meter settings for all individuals.

To assess the relationship between bill dimensions and bite force, we performed multiple regression analysis. We included bill depth and body size as predictors, because these traits have been found to strongly predict bite force in other passerines (Herrel et al., 2005; Soons et al., 2015; Van Der Meij and Bout, 2008). We generated a composite score of body size (PC1_bod_) using PCA of tarsometatarsus and wing lengths for individuals used in bite force analyses. We applied linear regression in R using the function *lm* with bill depth and PC1_bod_ as our predictors for maximum bite force. We did not include island sampled in our analyses as this was not independent of bill dimensions. We extracted standardized beta coefficients and 95% confidence intervals around unstandardized beta coefficients to compare the effects of both predictors on maximum bite force.

### Quantifying seed extraction time to infer the functional consequences of bill variation

We held a subset of captured males in 2014 for caged field trials to quantify foraging efficiency. Females were excluded from trials to prevent interruption of incubating or laying behaviors. Following capture, we immediately placed birds in covered trial cages for acclimation to experimental conditions (*see* Fig. S1 *for details*). All subjects were provided with water throughout the duration of the trial. We provided a two-hour acclimation and fasting period prior to the initiation of each trial. During the acclimation period, we monitored activity continuously via video cameras. Following acclimation, we initiated recording and slowly poured 10 grams of sterilized nyjer seed (*Guizotia abyssinica*) through a funnel in a brown plastic tube, which dispersed seeds across the floor of the cage. Sterilized nyjer seed is commonly used as bird feed for small passerines, and sterilization ensures the subsequent germination does not occur. Although *G. abyssinica* is not found on the Channel Islands and may not represent typical seed resources, song sparrows are generalist, ground-foragers. Thus, we assumed that our experimental food provisioning method facilitated normal foraging behavior. Trials lasted 45-120 minutes depending on latency to eat. We recorded behavioral notes during both acclimation and trial periods.

We reviewed foraging trial videos to quantify seed extraction time across multiple seeds. We counted the number of frames between when a bird’s bill tip lifted from the floor of the cage with a seed, to when the husk fell from the tomium (van der Meij and Bout, 2006). We divided the number of frames by the camera’s frame capture rate (29.97 frames/second) to calculate seed extraction time. The high temporal resolution of the cameras provided a precision of 0.033 seconds. We included only feeding events in which seed manipulation was observed throughout the entire seed extraction process.

Seed extraction is a complex task that requires manipulation of the bill along multiple axes. Consequently, we performed a PCA of bill depth, width, and length and used the first axis of variation (PC1_bill_) to test whether differences in bill morphology is result in differences in seed extraction time. We included PC1_bill_ of bill dimensions as a fixed effect in a linear mixed model predicting seed extraction time using the function *lmer* in the package *lme4* (Bates et al., 2015). We accounted for repeated observations of the same bird by including individual as a random effect. We extracted 95% confidence intervals around the estimated coefficient for PC1_bill_ using the function *confint* to infer the strength of PC1_bill_ in modifying seed extraction time.

## RESULTS

### Patterns of bill variation in contemporary populations

From 2014-2016, we measured 542 adult song sparrows (San Miguel Island, *n* = 104; Santa Rosa Island, *n* = 194; Santa Cruz Island, *n* = 244) sampled from 432 unique net locations (San Miguel Island, *n* = 68; Santa Rosa Island, *n* = 141; Santa Cruz Island, *n* = 223; Fig. 1). Patterns of bill variation in contemporary populations aligned with our expectations based on previous research using museum specimens (Greenberg and Danner 2012). Islands differed significantly in mean ranks for both raw bill surface area (*H*_df=2_ = 138.30, *P* < 0.001) and residual bill surface area (*H*_df=2_ = 143.96, *P* < 0.001) using Kruskal-Wallis tests. We found all Mann-Whitney U pairwise comparisons of raw and residual bill surface areas between islands were significant (Table 1). We confirmed larger bills were found on Santa Cruz Island, medium bills were found on Santa Rosa Island, and the smallest bills were found among San Miguel Island birds based on Hodges–Lehmann estimates of medians (Table 1).

**Table 1.**
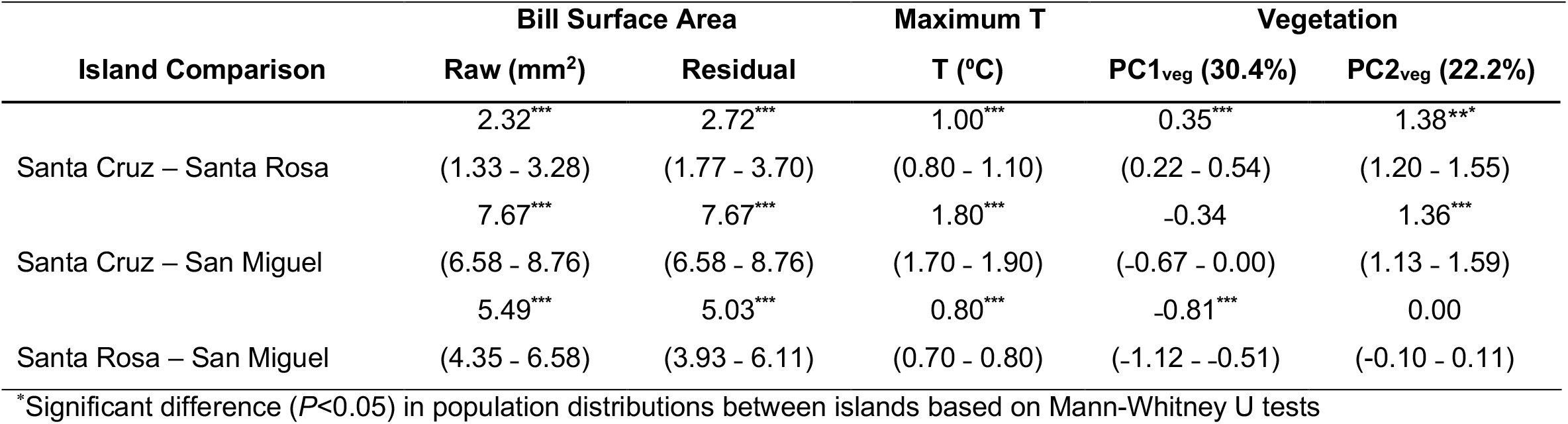
Nonparametric pairwise island comparisons (median±c.i.) for song sparrow raw (uncorrected for body size) and residual (corrected for body size) bill surface areas, maximum environmental temperatures (T), and vegetation dimensions (PC1, PC2). Pairwise comparisons were estimated using the Hodges-Lehmann method for bill surface areas in 542 birds (San Miguel Island, *n* = 104; Santa Rosa Island, *n* = 194; Santa Cruz Island, *n* = 244) and for temperature and vegetation characteristics across 432 unique sampling locations (San Miguel Island, *n* = 68; Santa Rosa Island, *n* = 141; Santa Cruz Island, *n* = 223). Vegetation dimensions include PC1 and PC2 from nonlinear PCA of ranked abundance in vegetation categories within breeding song sparrow territories. All islands had significantly different (*P* < 0.05) mean ranks in all bill and environmental variables based on nonparametric Kruskal-Wallis tests. Post-hoc Mann-Whitney U 95% confidence intervals around differences in mean ranks are shown in parentheses. Bill surface area analyses include only adult, breeding, territorial birds with complete phenotype measurements, and maximum temperature was extracted only for these unique sampling locations. Vegetation sampling occurred at most of these temperature sampling locations and at locations for territorial birds that did not have complete phenotype measurements.

| Island Comparison | Bill Surface Area |  | Maximum T | Vegetation |  |
| --- | --- | --- | --- | --- | --- |
|  | Raw (mm<sup>2</sup>) | Residual | T (°C) | PC1<sub>veg</sub> (30.4%) | PC2<sub>veg</sub> (22.2%) |
|  | 2.32*** | 2.72*** | 1.00*** | 0.35*** | 1.38*** |
| Santa Cruz – Santa Rosa | (1.33 - 3.28) | (1.77 - 3.70) | (0.80 - 1.10) | (0.22 - 0.54) | (1.20 - 1.55) |
|  | 7.67*** | 7.67*** | 1.80*** | -0.34 | 1.36*** |
| Santa Cruz – San Miguel | (6.58 - 8.76) | (6.58 - 8.76) | (1.70 - 1.90) | (-0.67 - 0.00) | (1.13 - 1.59) |
|  | 5.49*** | 5.03*** | 0.80*** | -0.81*** | 0.00 |
| Santa Rosa – San Miguel | (4.35 - 6.58) | (3.93 - 6.11) | (0.70 - 0.80) | (-1.12 - -0.51) | (-0.10 - 0.11) |

\*Significant difference ( $P < 0.05$ ) in population distributions between islands based on Mann-Whitney U tests

### Maximum temperature and vegetation within breeding territories across islands

Environmental conditions within song sparrow breeding territories differed among the 432 unique sampling locations across islands (San Miguel Island, *n* = 68; Santa Rosa Island, *n* = 141; Santa Cruz Island, *n* = 223; Fig. 1). We found mean ranks in island maximum temperatures were significantly different (*H*_2_ = 282.63, *P* < 0.001). Post-hoc pairwise comparisons of maximum temperature were significantly different between island pairs as expected, such that Santa Cruz Island had a higher and San Miguel has lower median estimates of maximum temperature in territories (Table 1). Additionally, we found ranked abundances in common vegetation types were significantly associated with island sampled using 10,000 Monte Carlo simulations in Fisher’s exact test (*P* < 0.001). Common dominant vegetation included coyote brush (*Baccharis pilularis*), toyon (*Heteromeles arbutifolia*), silver bush lupine (*Lupinus albifrons*), introduced sweet fennel (*Foeniculum vulgare*), and a mix of annual and perennial grasses (Table S1).

Using NLPCA, we reduced the complexity in correlated ranked abundance of vegetation types among sampling locations. The first two principal components explained a total of 52.7 % of the variation in vegetation. The first axis (PC1 _veg_) explained 30.5% of the variation in vegetation, and the abundance of grasses and the joint effects of the abundance of lupine, miscellaneous forbs, and other substrates (*i.e*., bare ground or rock, water) loaded in opposing directions (Fig. 2A). This suggests that positive values along PC1 _veg_ are indicative of territories with more lupine, forbs, and other substrates and less grass, and negative values represent the inverse of this relationship (Fig. 2A). The second axis of variation (PC2 _veg_) explained 22.2% of variation in vegetation and reflects a trade-off in fennel and coyote brush (Fig. 2A). Positive values indicate more fennel and less coyote brush, and negative values represent less fennel and more coyote brush (Fig. 2A). We found islands overlapped in vegetation space based on 95% kernel density contours (Fig. 2B). We assessed these relationships statistically using nonparametric tests and found significant differences among island mean ranks in both vegetation dimensions (PC1 _veg_, *H*_2_ = 31.73, *P* < 0.001; PC2 _veg_, *H*_2_ = 241.85, *P* < 0.001; Table 1). Using post-hoc Mann-Whitney U tests, we found PC1_veg_ scores for Santa Rosa Island territories were more negative, such that territories on Santa Rosa Island had more grass and less miscellaneous forbs, bare ground and rock, open water, and lupine compared to Santa Cruz and San Miguel Islands (Fig. 2B). In contrast, we found PC2 _veg_ scores for Santa Cruz Island territories were significantly greater, with Santa Cruz territories encompassing more fennel and less coyote brush compared to Santa Rosa and San Miguel Islands (Fig. 2B).

**Figure 2.**
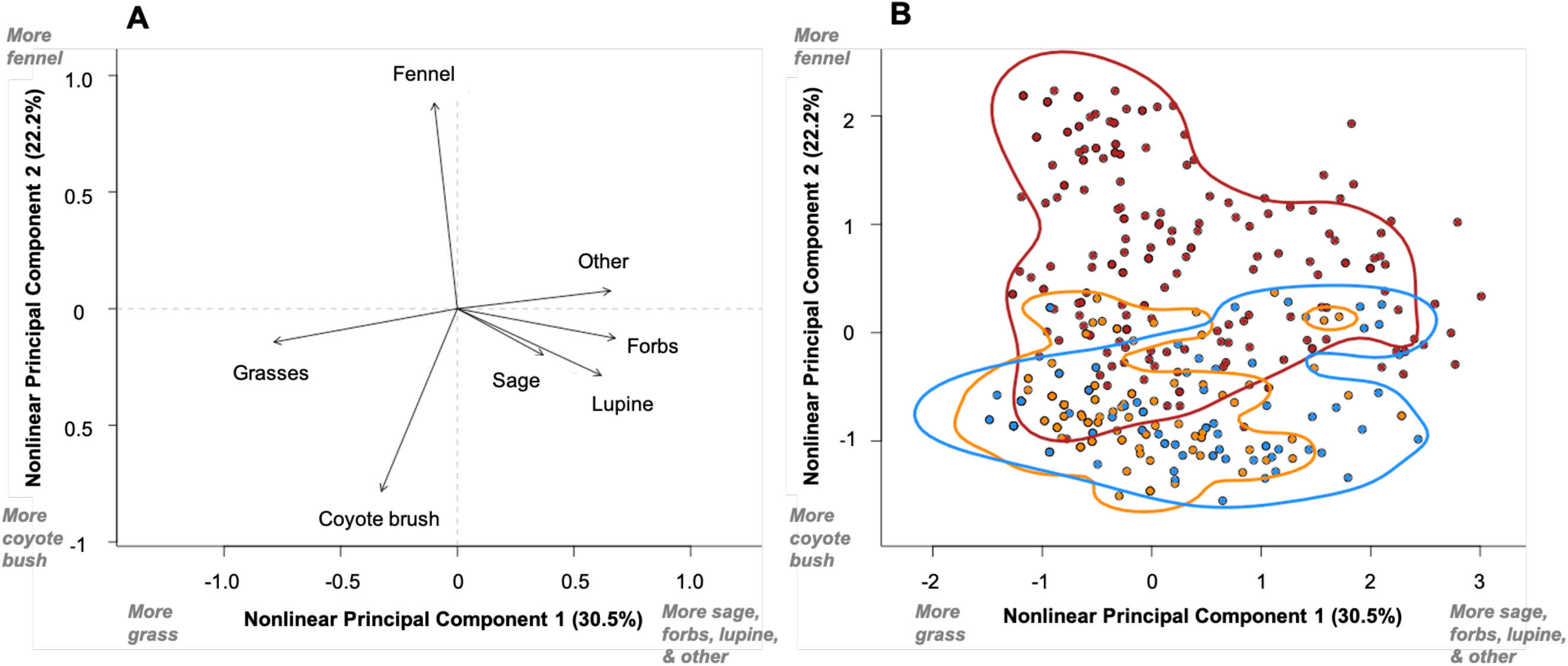
Variable loadings (A) and PC1, PC2, and 95% kernel density contours by island (B) from nonlinear PCA of vegetation within song sparrow territories. Sampling of 432 unique sampling locations occurred during the breeding season from 2014-2016 on San Miguel Island (blue; *n* = 68), Santa Rosa Island (orange; *n* = 141), and Santa Cruz Island (red; *n* = 223).

Multiple regression analysis was used to assess the relative importance of vegetation (PC1_veg_ and PC2_veg_) and maximum temperature in driving bill differences in adult song sparrows (Santa Cruz, *n* = 218 birds, Santa Rosa, *n* = 146 birds, San Miguel, *n* = 81 birds). These variables together explained a significant proportion of variation in residual bill surface area (*F*_3,442_ = 55.25, *p* < 0.001, adjusted R^2^ = 0.27; Fig. 3). We found residual bill surface area was significantly predicted by maximum island temperatures (β_standardized_ = 0.40, β_unstandardized_ = 3.06 (2.20 — 3.93), *t* = 10.350, *P* < 0.001). Although we found island differences in vegetation space, neither PC1_veg_ (β_standardized_ = −0.05, β_unstandardized_ = −0.28 (−0.77 — 0.20), *t* = −1.147, *P* = 0.25) nor PC2 _veg_ (β_standardized_ = 0.04, β_unstandardized_ = 0.20 (−0.43 — 0.83), *t* = 0.610, *P* = 0.54) were significant predictors. Vegetation space was used as a proxy for available food resources, and these results suggest climate, not food, is likely driving differences in residual bill surface area.

**Figure 3.**
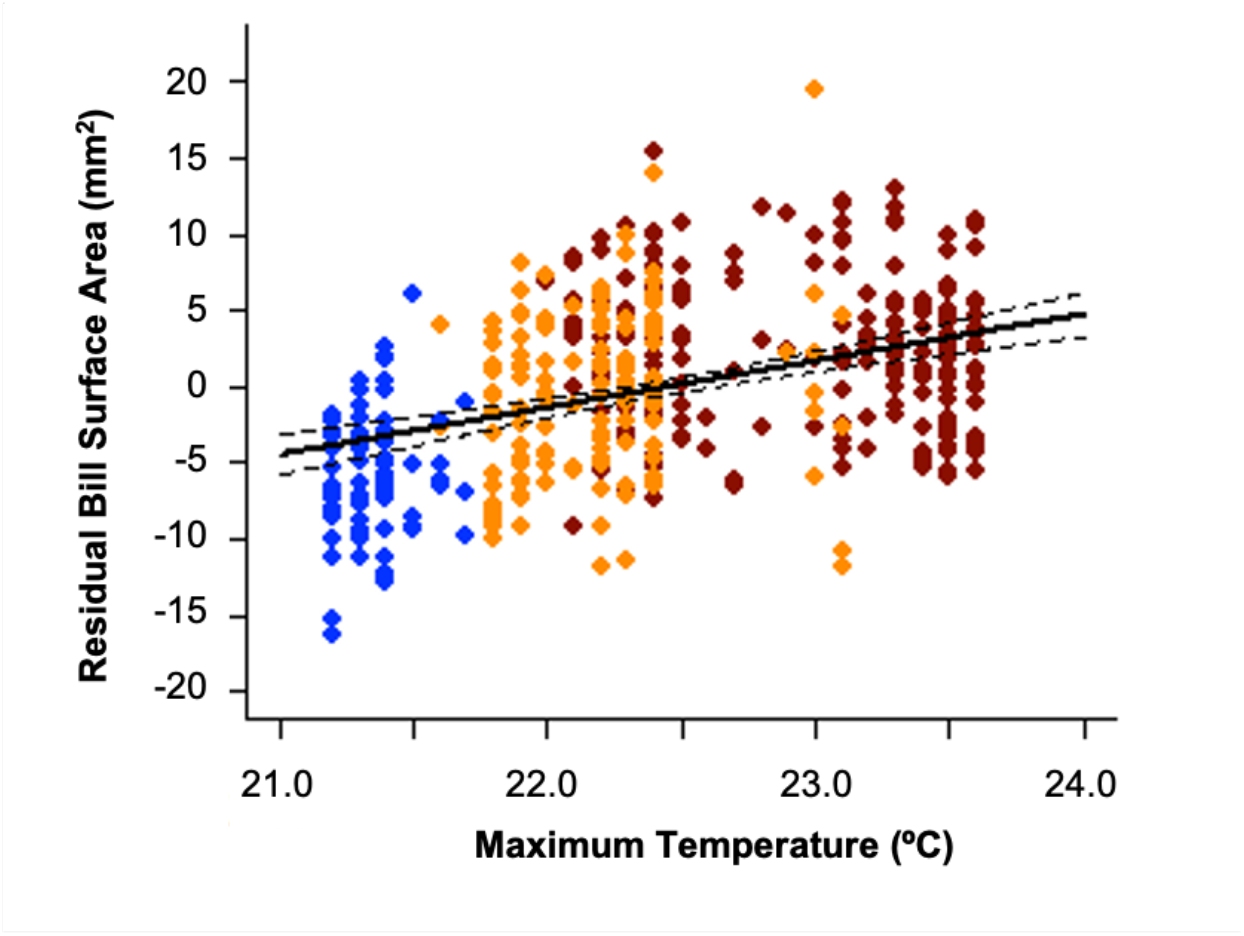
Residual bill surface area predicted by maximum temperature in song sparrows (*n* = 446) on the California Channel Islands. Residual bill surface area is total bill surface area corrected for skeletal body size and calculated from measures of bill depth, width, and length, tarsometatarsus length, and wing length in adult song sparrows on San Miguel Island (*n* = 81), Santa Rosa Island (orange, *n* = 147), and Santa Cruz Island (red, *n* = 218). Primary and secondary axes of variation from nonlinear PCA of vegetation (PC1_veg_ and PC2_veg_) were included in the linear regression analysis and were not significant predictors of residual bill surface area.

### Functional consequences of bill variation between island populations

Contrary to predictions from the foraging efficiency hypothesis, we did not find evidence that differences in bill morphology result in functional differences in bite force between birds on San Miguel (*n* = 28) and Santa Cruz Island (*n* = 28). Body size (PC1_bod_) and bill depth together explained very little of the variation in maximum bite force (*F*_2,53_ = 1.45, *P* = 0.24, adjusted R^2^ = 0.02; Fig. 4A). Although Santa Cruz Island birds tended to have larger structural bodies (PC_bod(Cruz)_, mean±sd = 0.22±1.09; PC_bod(Miguel)_, mean±sd = −0.22±1.02), body size distributions generally overlapped and did not significantly predict maximum bite force (β_standardized_ = 0.17, β_unstandardized_ = 0.23 (−0.14 — 0.59), *t* = 1.247, *P* = 0.22). Island sampling groups differed in bill depth as expected with birds on Santa Cruz Island having deeper bills (Depth_(Cruz)_, mean±sd = 6.15±0.21; Depth_(Miguel)_, mean±sd = 5.54±0.20; Fig. 4A). Yet, we found little evidence to suggest that these differences in depth result in synonymous changes in maximum bite force (β_standardized_ = 0.17, β_unstandardized_ = −0.74 (−1.80 — 0.32), *t* = −1.394, *P* = 0.17).

**Figure 4.**
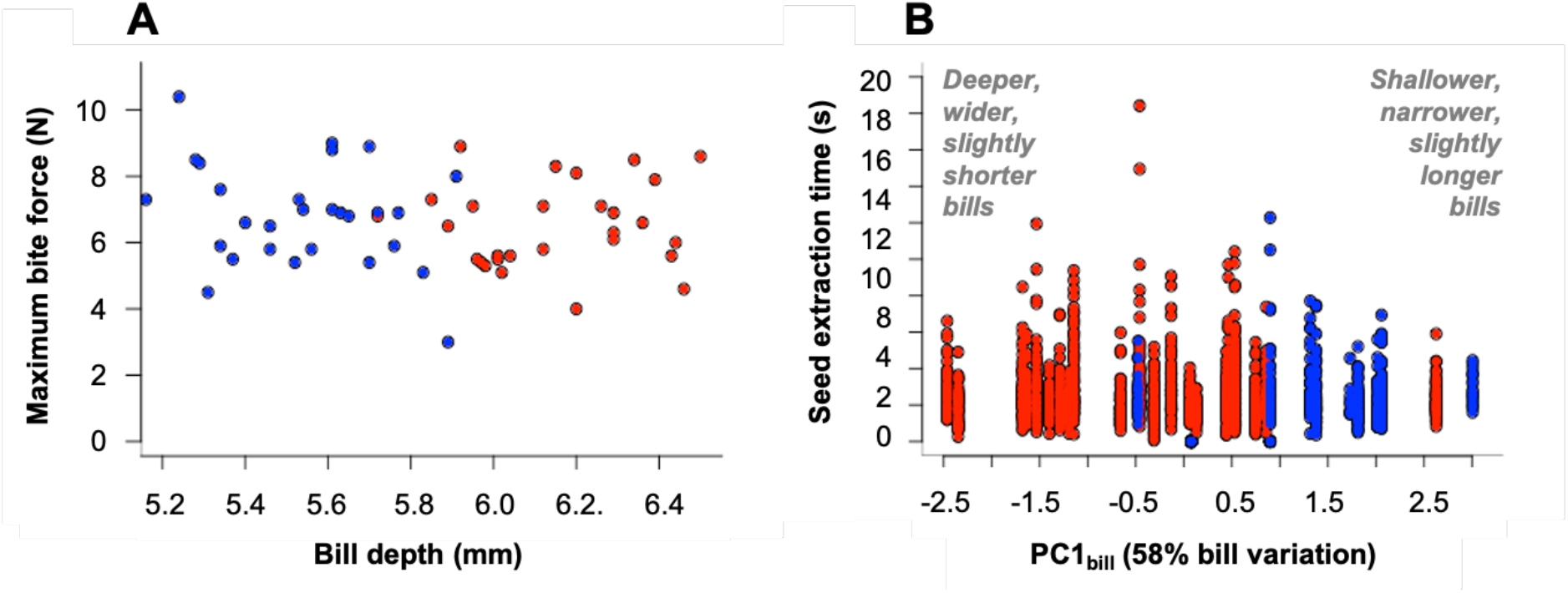
Relationship between song sparrow bill dimensions and foraging traits [maximum bite force (A) and seed extraction time (B)] between birds on Santa Cruz Island (red; *n*_A_ = 28, *n*_B_ = 23) and San Miguel Island (blue; *n*_A_ = 28, *n*_B_ = 10). PC1_bill_ is the first orthogonal axis in a PCA of bill depth, width, and length taken from the anterior edge of the nares.

We performed foraging trials in adult, male song sparrows from Santa Cruz Island (*n* = 23) and San Miguel Island (*n* = 10), and, again, found little evidence for an effect of bill morphology on seed extraction time. The first principal component (PC1_bill_) in a PCA of bill depth, width, and length explained 58% of the variation in bill dimensions and was largely driven by bill depth and width (Fig. S2). San Miguel Island and Santa Cruz Island birds overlapped along PC1 (Fig. 4B), but San Miguel birds loaded more positively (shallower, narrower, slightly longer bills). In contrast, Santa Cruz Island birds loaded more negatively, suggesting birds tended to have deeper, wider, slightly shorter bills. The mean number of observed extracted seeds was 60 seeds per individual (sd = 32, range = 8 – 163), and 13% of variance was explained by individual effects. PC1_bill_ had a very weak to negligible effect on seed extraction time (β = 0.009 (−0.14 – 0.16), *t*_32_ = 0.115, *P* = 0.91). These results combined with bite force analyses suggest that morphological differences in bills do not facilitate differences in foraging efficiency in island song sparrows.

## DISCUSSION

Identifying the ecological correlates of phenotypic variation provides insights into the selection pressures that may shape complex traits and their multiple functions. The avian bill is one such complex trait that has primarily been studied in the context of foraging ability despite its critical role in preening, communication, and thermoregulation. Indeed, there has been growing appreciation for the role of climate in shaping variation in bill morphology, as mounting evidence suggests the bill is an important tool for heat dissipation and thermoregulation (e.g., Gardner et al., 2016; Greenberg and Danner, 2013; Ryeland et al., 2017; Symonds and Tattersall, 2010). Here, we tested whether climate and/or vegetation composition within breeding territories significantly predicted bill variation in song sparrows on the California Channel Islands (i.e., San Miguel, Santa Rosa, and Santa Cruz Islands). We confirmed that bill size, maximum temperature, and vegetation composition differed among islands. However, only maximum temperature significantly predicted residual bill surface area in a multiple regression analysis including vegetation dimensions (PC1_veg_ and PC2_veg_) and temperature as predictors (Fig. 3). We did not find a significant relationship between bill morphology and either maximum bite force or seed extraction time. This provides additional evidence against foraging efficiency as a strong selective pressure in this system. Together, these results suggest that climate may be more important than diet and foraging ability in the evolution of a complex phenotype in song sparrows on the California Channel Islands.

### Vegetation differences do not directly result in concerted changes in bill morphology

Analysis of habitat composition with respect to vegetation provides key insights into the potential food resources available for breeding birds. Because song sparrows on the Channel Islands occupy a strong east-west climate gradient, we expected some degree of habitat differences among island breeding sites. Thus, it is not surprising that ranked abundance in our focal vegetation categories were significantly associated with island sampled. For example, Santa Cruz Island is characterized by greater heterogeneity in topography, soil composition, and climate, which is correlated with increases in species richness and diversity of plants compared to Santa Rosa and San Miguel Islands (Schoennerr et al., 2003). Our results demonstrate increased diversity in vegetation space (PC1_veg_ and PC2_veg_) within Santa Cruz Island territories (Fig. 2B). Yet, 95% kernel density contours also suggest a large proportion of overlap in vegetation space among islands (Fig. 2B), and similar plant taxa identifiable to the genus and species level were present across all islands (Table S1). More extensive vegetation and habitat sampling may allow us to parse out fine-scale habitat differentiation among islands, but whether any differences in vegetation result in differences in food resources is uncertain. Direct observation of foraging behavior and evaluation of the available dietary resources during the non-breeding season, when song sparrows most heavily rely on seeds, would allow us to draw stronger inferences regarding the relationship among foraging ability, diet, and bill morphology. Nevertheless, our analyses support the hypothesis that climate may facilitate vegetative differences among song sparrow territories, but differences in vegetation did not directly predict bill morphology in multiple regression analyses. Hence, vegetation, a proxy for food resources and diet, may not be a strong selective pressure generating bill variation across islands, and we found further evidence against selection operating on foraging ability in experimental tests of foraging efficiency (Figs. 4A-B).

### Variation in bill morphology does not result in differences in foraging efficiency

We did not find a relationship between variation in bill morphology and either bite force or seed extraction time (Fig. 4A-B), despite evidence that bill dimensions affect feeding performance in other passerines (Anderson et al., 2008; Herrel et al., 2005; Navalón et al., 2019). Indeed, some of the most well-documented cases of specialization in resource acquisition with respect to bill variation occur in island systems (Burns et al., 2003; Grant and Grant, 2002). Of these cases, Darwin’s ground finches (*Geospiza sp*.) are perhaps most notable and ecologically similar to song sparrows in their foraging behavior and diet (De León et al., 2014). In Darwin’s finches, birds exhibit correlations between both bill dimensions and bite force (Herrel et al., 2005; van der Meij and Bout, 2008; Soons et al., 2015) and bill dimensions and seed extraction times (van der Meij and Bout, 2008). The discordance between our results and findings from studies of Darwin’s finches may be attributed to multiple factors.

First, sparrows are generalist foragers with a heavy insectivorous diet during the breeding season and a transition to a granivorous diet during the non-breeding season (Arcese et al., 2002), whereas Darwin’s ground finches’ diets consist primarily of seeds throughout the year (De León et al., 2014). Specialization on seeds and food limitation in ground finches facilitate competitive interactions among species that overlap in their dietary niches (De León et al., 2014), resulting in strong selection on individuals to optimize bill morphology for increased foraging efficiency. On the California Channel Islands, it is unclear whether food resources are limited and whether food limitation imposes a strong selective pressure on song sparrow populations. Mass comparisons between sparrows from the Channel Islands and nearby mainland California found island sparrows to heavier than mainland sparrows when correcting for structural body size, suggesting island birds may be in better condition (Danner et al., 2014). Increased mass among island sparrows supports the hypotheses that food resources are not limited and reduced interspecific competition for food favors the sparrows’ generalist diets (Blondel, 2000; Clegg, 2010; Diamond, 1970; Keast, 1970; Scott et al., 2003). As a result, variation in bill morphology may be decoupled from foraging efficiency traits measured in this study.

Additionally, methodological limitations may have prevented us from quantifying key traits associated with foraging. For example, previous research in Darwin’s finches identify a link between muscle mass and maximum bite force (Herrel et al., 2005; Herrel et al., 2010; Soons et al., 2015), and other skeletal features are correlated with bite force and closing velocity (Corbin et al., 2015). These methods require analysis of euthanized individuals, which we were unable perform. Here, we evaluated bite force differences elicited from the posterior section of the bill, not from the anterior (the tip) where functional differences in grasping and object manipulation occur across avian families (Clayton et al., 2005; Demery et al., 2011; Gentle et al., 1982; Sustaita, 2007). Although skeletal structures associated with foraging traits are highly-correlated (Van Der Meij and Bout, 2008) and our measures adequately estimate a large proportion of variation in overall bill morphology, future studies may examine unmeasured phenotypic traits in this study to better estimate the functional consequences of craniofacial variation. Finally, we were limited to using readily available bird seed for foraging trials to increase our ability to observe detailed manipulation of the food resource. We did not compare nyjer seed characteristics with those used by song sparrows on the Channel Islands during the winter months. Yet, seed availability during winter months were likely similar given that non-native, seed-producing plant species (e.g., annual grasses) are widespread (Junak et al., 2007) and were present at most sampling locations (Table S1). Sample size does not appear to be an issue, as a power analysis suggests that our sample sizes were sufficient to detect a biologically relevant difference in maximum bite force given our experimental design (Fig. S3). Although we did not find evidence that foraging efficiency is an important driver of variation in bill shape in our study populations, further research is needed to definitively exclude the possibility that differing food resources among islands is a selective force. Our experimental design provides a framework for future studies to test how bill dimensions influence multiple components of foraging efficiency (i.e., both bite force and seed extraction).

### Evidence for the bill as a thermoregulatory trait in island sparrows

Our results are consistent with the hypothesis that selection operates on the bill to improve thermoregulatory ability in passerines occupying xeric environments. Increasing empirical evidence demonstrate a relationship between climate and bill morphology that aligns with the thermoregulatory hypothesis (reviewed by Tattersall et al., 2016), and this relationship may be traced over evolutionary time (Campbell-Tennant et al., 2015). The ability to radiate heat from unfeathered structures is particularly important for small passerines, including song sparrows, which are more vulnerable to dehydration from evaporative water loss and, thus, more susceptible to adverse effects of thermal stress (Mckechnie and Wolf, 2010; Whitfield et al., 2015). Indeed, our results are consistent with previous research that identified a significant, positive correlation between climate and bill morphology in eastern and Atlantic song sparrows (Danner and Greenberg, 2014). A similar pattern has been described in Darwin’s finches (Tattersall et al., 2018) and other similarly-sized passerines (Greenberg and Danner, 2012; LaBarbera et al., 2017; Laiolo and Rolando, 2001). The magnitude of the effect of climate in predicting bill morphology may change according to seasonality (Greenberg et al., 2013), environmental variation during development (Burness et al., 2013; Labarbera et al., 2020), habitat type (Luther and Greenberg, 2014), and sex (Greenberg and Danner, 2013). Importantly, selection may act simultaneously on other traits to facilitate thermoregulation, including internal nasal structures (Danner et al., 2017), plumage (Wolf and Walsberg, 2000), and physiological performance (e.g., Noakes et al., 2016; Tieleman et al., 2003; White et al., 2007; Whitfield et al., 2015). Thus, further research that explores the complex relationship between temperature, humidity, and other morphological and physiological traits is needed to better understand how climate facilitates and maintains phenotypic variation.

### Conclusions

Consistent with previous studies using museum specimens, variation in bill morphology among contemporary song sparrow populations on the California Channel Island is correlated with maximum temperature, suggesting an important thermoregulatory function. Differences in the vegetation and habitats used by sparrows on different islands were not strongly predictive of observed bill divergence. Variation in bill morphology was also not correlated with bite force or seed extraction, perhaps because song sparrows are generalist foragers. We hope that our results encourage future research about how different environmental agents of selection simultaneously act on avian bills to optimize the multiple, fitness-related functions of foraging, thermoregulation, preening, and vocalization.

## ACKNOWLEDGEMENTS

We dedicate this paper to our late colleague Dr. Russ Greenberg (1953-2013), who played a formative role in study design for this project. We acknowledge, honor, and respect the people of the Chumash Nation, the original stewards of coastal southern California and the Northern California Channel Islands [San Miguel (*Tuqan*), Santa Rosa (*Wi’ma*), Santa Cruz (*Limuw*), and Anacapa Islands (*Anyapakh*)]. We are grateful to the many agencies and individuals that aided in the collection of data through transportation and field support including The Nature Conservancy in California (Jennifer Baker, Christina Boser, David Dewey, and Eamon O’Byrne), UCSB Santa Cruz Island Reserve staff (Lyndall Laughrin, Lynn McLaren, and Brian Guerrero), Channel Islands National Park Service (Linda Dye, Tim Coonan, Ian Williams, Lulis Cuevas, Paula Power, David Mazurkiewicz, Kate Faulkner and Ken Convery), CSUCI Santa Rosa Island Research Station staff (Cause, Tracy, and Solstice Hanna, Robyn Shea, Aspen Coty, and Russell Bradley), and field technicians (Janelle Chojnacki, Carolyn Cummins, Graham Sorenson, Eamon Harrity, Angela Hsiung, and Kathryn Fleming).

## AUTHOR CONTRIBUTIONS

R.M.D., J.F.H., and T.S.S. designed the study. R.M.D. and M.P.G. collected field data with additional support from T.S.S. J.F.H. provided instrumentation for field data collection. R.A.F. extracted foraging data from videos. M.P.G. conducted all analyses with support from R.M.D. The article was written by M.P.G. with additional input from C.A.M., W.C.F., T.S.S., J.F.H., and R.M.D.

## DATA AVAILABILITY

Phenotypic and environmental data used for all analyses are available on Dryad (https://doi.org/10.5061/dryad.wwpzgmsjc)

## LIST OF SYMBOLS AND ABBREVIATIONS

PC1_bod_: body size; composite score of body size based on PCA of tarsometatarsus and wing lengths used to estimate residual bill surface area
PC1_veg_: primary axis of variation in vegetation; composite score of vegetation based on NLPCA of common vegetation types in sampling locations
PC2_veg_: secondary axis of variation in vegetation; composite score of vegetation based on NLPCA of common vegetation types in sampling locations
PC1_bill_: primary axis of variation in bill dimensions; composite score of bill depth, width, and length used in analysis of foraging efficiency
T: maximum environmental temperature (in July) of sampling location

## REFERENCES

Åkesson, M., Bensch, S., Hasselquist, D., Tarka, M. and Hansson, B. (2008). Estimating heritabilities and genetic correlations: comparing the “animal mode” with parent-offspring regression using data from a natural population. PLoS One 3, e1739.

Anderson, R. A., Mcbrayer, L. D. and Herrel, A. (2008). Bite force in vertebrates: Opportunities and caveats for use of a nonpareil whole-animal performance measure. Biol. J. Linn. Soc. 93, 709–720.

Arcese, P., Sogge, M. K., Marra, A. B. and Patten, M. A. (2002). Song sparrow (Melospiza meldoia). Birds North Am. Online.

Ballentine, B. (2006). Morphological adaptation influences the evolution of a mating signal. Evolution 60, 1936–44.

Barbosa, A. and Moreno, E. (1999). Evolution of foraging strategies in shorebirds: an ecomorphological approach. Auk 116, 712–725.

Bates, D., Mächler, M., Bolker, B. M. and Walker, S. C. (2015). Fitting linear mixed-effects models using lme4. J. Stat. Softw. 67, 1–48.

Behrendt, S. (2014). lm.beta: add standardized regression coefficients to lm-objects.

Benkman, C. W. (1993). Adaptation to single resources and the evolution of crossbill (Loxia) diversity. Ecol. Monogr. 63, 305–325.

Benkman, C. W. (2003). Divergent selection drives the adaptive radiation of crossbills. Evolution 57, 1176–1181.

Blondel, J. (2000). Evolution and ecology of birds on islands: trends and prospects. Vie Milieu 50, 205–220.

Boag, P. T. (1983). The heritability of external morphology in Darwin’s ground finches (Geospiza) on Isla Daphne Major, Galapagos. Evolution 37, 877–894.

Burness, G., Huard, J. R., Malcolm, E. and Tattersall, G. J. (2013). Post-hatch heat warms adult beaks: irreversible physiological plasticity in Japanese quail. Proc. R. Soc. B Biol. Sci. 280, 20131436.

Burns, K. J., Hackett, S. J. and Klein, N. K. (2003). Phylogenetic relationships of neotropical honeycreepers evolution of feeding morphology. J. Avian Biol. 34, 360–370.

Campbell-Tennant, D. J. E., Gardner, J. L., Kearney, M. R. and Symonds, M. R. E. (2015). Climate-related spatial and temporal variation in bill morphology over the past century in Australian parrots. J. Biogeogr. 42, 1163–1175.

Clayton, D. H., Moyer, B. R., Bush, S. E., Jones, T. G., Gardiner, D. W., Rhodes, B. B. and Goller, F. (2005). Adaptive significance of avian beak morphology for ectoparasite control. Proc. Biol. Sci. 272, 811–817.

Clegg, S. M. (2010). Evolutionary changes following island colonization in birds: empirical insights into the roles of microevolutionary processes. In The theory of island biogeography revisited (ed. Losos, J. B.) and Ricklefs, R. E.), pp. 293–325. Princeton, New Jersey: Princeton University Press.

Clinchy, M., Sheriff, M. J. and Zanette, L. Y. (2013). Predator-induced stress and the ecology of fear. Funct. Ecol. 27, 56–65.

Corbin, C. E., Lowenberger, L. K. and Gray, B. L. (2015). Linkage and trade-off in trophic morphology and behavioural performance of birds. Funct. Ecol. 29, 808–815.

Danner, R. M. and Greenberg, R. (2014). A critical season approach to Allen’s rule: Bill size declines with winter temperature in a cold temperate environment. J. Biogeogr. 114–120.

Danner, R. M., Greenberg, R. and Sillett, T. S. (2014). The implications of increased body size in the song sparrows of the California Islands. Monogr. West. North Am. Nat. 7, 348–356.

Danner, R. M., Gulson-castillo, E. R., James, H. F., Dzielski, S. A., Iii, D. C. F., Sibbald, E. T. and Winkler, D. W. (2017). Habitat-specific divergence of air conditioning structures in bird bills. Auk 134, 65–75.

Dawson, W. R. (1981). Evaporative losses of water by birds. Comp. Biochem. Physiol. A. Mol. Integr. Physiol. 71A, 495–509.

De León, L. F., Podos, J., Gardezi, T., Herrel, A. and Hendry, A. P. (2014). Darwin’s finches and their diet niches: The sympatric coexistence of imperfect generalists. J. Evol. Biol. 27, 1093–1104.

Demery, Z. P., Chappell, J. and Martin, G. R. (2011). Vision, touch and object manipulation in senegal parrots Poicephalus senegalus. Proc. R. Soc. B Biol. Sci. 278, 3687–3693.

Diamond, J. M. (1970). Ecological consequences of island colonization by southwest Pacific birds, I. Types of niche shifts. Proc. Natl. Acad. Sci. 67, 529–536.

Egea-Serrano, A., Hangartner, S., Laurila, A. and Räsänen, K. (2014). Multifarious selection through environmental change: Acidity and predator-mediated adaptive divergence in the moor frog (Rana arvalis). Proc. R. Soc. B Biol. Sci. 281, 20133266.

Fayet, A. L., Hansen, E. S. and Biro, D. (2020). Evidence of tool use in a seabird. Proc. Natl. Acad. Sci. U. S. A. 117, 1277–1279.

Friedman, N. R., Miller, E. T., Ball, J. R., Kasuga, H., Remeš, V. and Economo, E. P. (2019). Evolution of a multifunctional trait: Shared effects of foraging ecology and thermoregulation on beak morphology, with consequences for song evolution. Proc. R. Soc. B Biol. Sci. 286,.

Gardner, J. L., Symonds, M. R. E., Joseph, L., Ikin, K., Stein, J. and Kruuk, L. E. B. (2016). Spatial variation in avian bill size is associated with humidity in summer among Australian passerines. Clim. Chang. Responses 3, 1–11.

Gentle, M. J., Hughes, B. O. and Hubrecht, R. C. (1982). The effect of beak trimming on food intake, feeding behaviour and body weight in adult hens. Appl. Anim. Ethol. 8, 147–159.

Ghalambor, C. K., Walker, J. A. and Reznick, D. N. (2003). Multi-trait selection, adaptation, and constraints on the evolution of burst swimming performance. Integr. Comp. Biol. 438, 431–438.

Grant, P. R. (1983). Inheritance of size and shape in a population of Darwin’s finches, Geospiza conirostris. Proc. R. Soc. London - Biol. Sci. 220, 219–236.

Grant, P. R. and Grant, B. R. (2002). Adaptive radiation of Darwin’s finches. Am. Sci. 90, 130–139.

Greenberg, R. and Danner, R. M. (2012). The influence of the California marine layer on bill size in a generalist songbird. Evolution 66, 3825–35.

Greenberg, R. and Danner, R. M. (2013). Climate, ecological release and bill dimorphism in an island songbird. Biol. Lett. 9, 20130118.

Greenberg, R., Cadena, V., Danner, R. M. and Tattersall, G. J. (2012). Heat loss may explain bill size differences between birds occupying different habitats. PLoS One 7, e40933.

Greenberg, R., Etterson, M. and Danner, R. M. (2013). Seasonal dimorphism in the horny bills of sparrows. Ecol. Evol. 3, 389–98.

Hagan, A. and Heath, J. E. (1980). Regulation of heat loss in the duck by vasomotion in the bill. J. Therm. Biol. 5, 95–101.

Herrel, A., Spithoven, L., Van Damme, R. and De Vree, F. (1999). Sexual dimorphism of head size in Gallotia galloti: Testing the niche divergence hypothesis by functional analyses. Funct. Ecol. 13, 289–297.

Herrel, A., Podos, J., Huber, S. K. and Hendry, A. P. (2005). Evolution of bite force in Darwin’s finches: A key role for head width. J. Evol. Biol. 18, 669–675.

Herrel, A., Soons, J., Aerts, P., Dirckx, J., Boone, M., Jacobs, P., Adriaens, D. and Podos, J. (2010). Adaptation and function of the bills of Darwins finches: Divergence by feeding type and sex. Emu 110, 39–47.

Hijmans, R. J., Cameron, S. E., Parra, J. L., Jones, P. G. and Jarvis, A. (2005). Very high resolution interpolated climate surfaces for global land areas. Int. J. Climatol. 25, 1965–1978.

Jensen, H., Saether, B.-E., Ringsby, T. H., Tufto, J., Griffith, S. C. and Ellegren, H. (2003). Sexual variation in heritability and genetic correlations of morphological traits in house sparrow (Passer domesticus). J. Evol. Biol. 16, 1296–1307.

Jones, J. S., Leith, B. H. and Rawlings, P. (1977). Polymorphism in Cepaea: A problem with too many solutions? Annu. Rev. Ecol. Syst. 8, 109–143.

Junak, S., Knapp, D. A., Haller, J. R., Philbrick, R., Schoenherr, A. and Keeler-Wolf, T. (2007). The California Channel Islands. In Terrestrial Vegetation of California, p. University of California Press.

Kawecki, T. J. and Ebert, D. (2004). Conceptual issues in local adaptation. Ecol. Lett. 7, 1225–1241.

Keast, A. (1970). Adaptive evolution and shifts in niche occupation in island birds. Assoc. Trop. Biol. Conserv. 2, 61–75.

Keller, L. F., Grant, P. R., Rosemary Grant, B. and Petren, K. (2001). Heritability of morphological traits in Darwin’s finches: Misidentified paternity and maternal effects. Heredity 87, 325–336.

Kim, S. Y., Noguera, J. C., Morales, J. and Velando, A. (2011). Quantitative genetic evidence for trade-off between growth and resistance to oxidative stress in a wild bird. Evol. Ecol. 25, 461–472.

Labarbera, K., Marsh, K. J., Hayes, K. R. R. and Hammond, T. T. (2020). Context-dependent effects of relative temperature extremes on bill morphology in a songbird. R. Soc. Open Sci. 7, 192203

LaBarbera, K., Hayes, K. R., Marsh, K. J. and Lacey, E. A. (2017). Complex relationships among environmental conditions and bill morphology in a generalist songbird. Evol. Ecol. 31, 707–724.

Laiolo, P. and Rolando, A. (2001). Ecogeographic correlates of morphometric variation in the red-billed Chough Pyrrhocorax pyrrhocorax and the Alpine Chough Pyrrhocorax graculus. Ibis 143, 602–616.

Lamichhaney, S., Berglund, J., Almén, M. S., Maqbool, K., Grabherr, M., Martinez-Barrio, A., Promerová, M., Rubin, C.-J., Wang, C., Zamani, N., et al. (2015). Evolution of Darwin’s finches and their beaks revealed by genome sequencing. Nature 518, 371–375.

Langin, K. M., Sillett, T. S., Funk, W. C., Morrison, S. A., Desrosiers, M. A. and Ghalambor, C. K. (2015). Islands within an island: Repeated adaptive divergence in a single population. Evolution (N. Y). 69, 653–665.

Lerner, H. R. L., Meyer, M., James, H. F., Hofreiter, M. and Fleischer, R. C. (2011). Multilocus resolution of phylogeny and timescale in the extant adaptive radiation of Hawaiian honeycreepers. Curr. Biol. 21, 1838–1844.

Lima, S. L. and Dill, L. M. (1990). Behavioral decisions made under the risk of predation: A review and prospectus. Can. J. Zool. 68, 619–640.

Luther, D. and Greenberg, R. (2014). Habitat type and ambient temperature contribute to bill morphology. Ecol. Evol. 4, 699–705.

MacColl, A. D. C. (2011). The ecological causes of evolution. Trends Ecol. Evol. 26, 514–522.

Mair, P. and de Leeuw, J. (2019). Gifi: Multivariate Analysis with Optimal Scaling.

Mckechnie, A. E. and Wolf, B. O. (2010). Climate change increases the likelihood of catastrophic avian mortality events during extreme heat waves. Biol. Lett. 6, 253–256.

Navalón, G., Bright, J. A., Marugán-Lobón, J. and Rayfield, E. J. (2019). The evolutionary relationship among beak shape, mechanical advantage, and feeding ecology in modern birds. Evolution 73, 422–435.

Nebel, S., Jackson, D. L. and Elner, R. W. (2005). Functional association of bill morphology and foraging behaviour in calidrid sandpipers. Anim. Biol. 55, 235–243.

Noakes, M., Wolf, B. and McKechnie, A. (2016). Seasonal and geographical variation in heat tolerance and evaporative cooling capacity in a passerine bird. J. Exp. Biol. 219, 859–869.

Parchman, T. L., Benkman, C. W. and Britch, S. C. (2006). Patterns of genetic variation in the adaptive radiation of New World crossbills (Aves: Loxia). Mol. Ecol. 15, 1873–1887.

Pfrender, M. E. (2012). Triangulating the genetic basis of adaptation to multifarious selection. Mol. Ecol. 21, 2051–2053.

Podos, J. (2001). Correlated evolution of morphology and vocal signal structure in Darwin’s finches. Nature 409, 185–188.

Podos, J. and Nowicki, S. (2004). Beaks, adaptation, and vocal evolution in Darwin’s finches. Bioscience 54, 501.

Reznick, D. N. and Travis, J. (1996). The empirical study of adaptations in natural populations. In Adaptations (ed. Rose, M. R.) and Lauder, G. V.), pp. 243–290. San Diego, California: Academic Press.

Rising, J. D. and Somers, K. M. (1989). The measurement of overall body size in birds. Auk 106, 666–674.

Robinson, M. R., Pilkington, J. G., Clutton-Brock, T. H., Pemberton, J. M. and Kruuk, L. E. B. (2006). Live fast, die young: Trade-offs between fitness components and sexually antagonistic selection on weaponry in soay sheep. Evolution 60, 2168.

Rutz, C., Klump, B. C., Komarczyk, L., Leighton, R., Kramer, J., Wischnewski, S., Sugasawa, S., Morrissey, M. B., James, R., St Clair, J. J. H., et al. (2016). Discovery of species-wide tool use in the Hawaiian crow. Nature 537, 403–407.

Ryeland, J., Weston, M. A. and Symonds, M. R. E. (2017). Bill size mediates behavioural thermoregulation in birds. Funct. Ecol. 31, 885–893.

Schoennerr, A. A., Feldmeth, C. R. and Emerson, M. J. (2003). The Natural History of the Islands of California. Berkeley: University of California Press.

Scott, S. N., Clegg, S. M., Blomberg, S. P., Kikkawa, J. and Owens, I. P. F. (2003). Morphological shifts in island-dwelling birds: The roles of generalist foraging and niche expansion. Evolution 57, 2147–2156.

Shuford, W. D. and Gardali, T. eds. (2008). Channel Island song sparrow. In California bird species of special concern: A ranked assessment of species, subspecies, and distinct populations of birds of immediate conservation concern in California. Studies of Western Birds 1., p. Camarillo, CA: Western Field Ornithologists and California Department of Fish and Game.

Shultz, A. J. and Burns, K. J. (2017). The role of sexual and natural selection in shaping patterns of sexual dichromatism in the largest family of songbirds (Aves: Thraupidae). Evolution 71, 1061–1074.

Siepielski, A. M., DiBattista, J. D. and Carlson, S. M. (2009). It’s about time: The temporal dynamics of phenotypic selection in the wild. Ecol. Lett. 12, 1261–76.

Siepielski, A. M., Gotanda, K. M., Morrissey, M. B., Diamond, S. E., DiBattista, J. D. and Carlson, S. M. (2013). The spatial patterns of directional phenotypic selection. Ecol. Lett. 16, 1382–92.

Soons, J., Genbrugge, A., Podos, J., Adriaens, D., Aerts, P., Dirckx, J. and Herrel, A. (2015). Is beak morphology in Darwin’s finches tuned to loading demands? PLoS One 10,.

Sustaita, D. (2007). Musculoskeletal underpinnings to differences in killing behavior between North American accipiters (Falconiformes: Accipitridae) and falcons (Falconidae). J. Morphol. 269, 283–301.

Svensson, E. I. and Calsbeek, R. eds. (2012). The adaptive landscape in evolutionary biology. Oxford, United Kingdom: Oxford University Press.

Symonds, M. R. E. and Tattersall, G. J. (2010). Geographical variation in bill size across bird species provides evidence for Allen’s rule. Am. Nat. 176, 188–97.

Tattersall, G. J., Andrade, D. V and Abe, A. S. (2009). Heat exchange from the toucan bill reveals a controllable vascular thermal radiator. Science 325, 468–70.

Tattersall, G. J., Arnaout, B. and Symonds, M. R. E. (2016). The evolution of the avian bill as a thermoregulatory organ. Biol. Rev.

Tattersall, G. J., Chaves, J. A. and Danner, R. M. (2018). Thermoregulatory windows in Darwin’s finches. Funct. Ecol. 32, 358–368.

Temeles, E. J. and Kress, W. J. (2003). Adaptation in a plant-hummingbird association. Science 300, 630–633.

Temeles, E. J., Roberts, W. M., Url, S. and Mark, W. (1993). Dimorphism in bill length on foraging behavior: An experimental analysis of hummingbirds. Oecologia 94, 87–94.

Templeton, C. N. and Shriner, W. M. (2004). Multiple selection pressures influence Trinidadian guppy (Poecilia reticulata) antipredator behavior. Behav. Ecol. 15, 673–678.

Tieleman, B. I., Williams, J. B., Buschur, M. E. and Brown, C. R. (2003). Phenotypic variation of larks along an aridity gradient: Are desert birds more flexible? Ecology 84, 1800–1815.

Troscianko, J., Von Bayern, A. M. P., Chappell, J., Rutz, C. and Martin, G. R. (2012). Extreme binocular vision and a straight bill facilitate tool use in New Caledonian crows. Nat. Commun. 3, 1–7.

Van Der Meij, M. A. A. and Bout, R. G. (2004). Scaling of jaw muscle size and maximal bite force in finches. J. Exp. Biol. 207, 2745–2753.

Van Der Meij, M. A. A. and Bout, R. G. (2006). Seed husking time and maximal bite force in finches. J. Exp. Biol. 209, 3329–3335.

Van Der Meij, M. A. A. and Bout, R. G. (2008). The relationship between shape of the skull and bite force in finches. J. Exp. Biol. 211, 1668–1680.

White, C. R., Blackburn, T. M., Martin, G. R. and Butler, P. J. (2007). Basal metabolic rate of birds is associated with habitat temperature and precipitation, not primary productivity. Proc. Biol. Sci. 274, 287–93.

Whitfield, M. C., Smit, B., McKechnie, A. E. and Wolf, B. O. (2015). Avian thermoregulation in the heat: scaling of heat tolerance and evaporative cooling capacity in three southern African arid-zone passerines. J. Exp. Biol. 218, 1705–1714.

Wilkins, M. R., Seddon, N. and Safran, R. J. (2013). Evolutionary divergence in acoustic signals: Causes and consequences. Trends Ecol. Evol. 28, 156–166.

Wolf, B. O. and Walsberg, G. E. (2000). The role of the plumage in heat transfer processes of birds. Am. Zool. 40, 575–584.

