## Supplemental Materials for "Environmental correlates and functional consequences of bill divergence in island song sparrows"

**Table S1. Vegetation presence (x) within song sparrow territories on three California Channel Islands [San Miguel Island (*n* = 58), Santa Rosa Island (*n* = 112), and Santa Cruz Island (*n* = 183)].**

**Figure S1.** Experimental chambers for in-situ foraging efficiency trials conducted on song sparrows from San Miguel and Santa Cruz Islands during the breeding season (Feb-June) in 2014. Trial cages (44.5 cm x 22.2 cm x 26.7 cm) were equipped with two perches at different heights and absorbent material to collect excess fecal matter and minimize stress. We provided a plastic, visual block on one of the four sides of each cage to protect subjects from potential predators and extreme weather conditions which may influence stress and willingness to forage (Clinchy et al., 2013; Lima and Dill, 1990). We secured a translucent cloth on three of the remaining four sides to allow for light to enter thereby stimulating activity. The plastic shield contained a hole fitted to the lens of high definition Sanyo Dual Camera Xacti WH1 camcorders (Sanyo Model VPC-WH1YL), which were mounted on the outer side of the shield on flexible tripods.

**Figure S2. Variable loadings of principal component analyses (PCA) of bill dimensions [depth (billd), length (billl), and width (billw)] for analyses of foraging efficiency.** PCA was conducted on bill dimensions in 33 individuals (San Miguel Island, *n* = 10; Santa Cruz Island, *n* = 23) to generate a composite score of overall bill morphology. Bill depth and width loaded equally (44%), and bill length did not strongly load on PC1 (12%).

**Figure S3. Results from power analyses for bite force.** We performed power analyses to test if sample sizes were sufficient to detect biologically relevant differences between paired populations. We predicted biologically relevant differences in bite force would be 1.5 Newtons. We simulated data based on our means and variances measured in this study and then compared the model fit following Bolker (2009).

**Table S1. Vegetation presence (x) within song sparrow territories on three California Channel Islands [San Miguel Island ($n$ = 58), Santa Rosa Island ($n$ = 112), and Santa Cruz Island ($n$ = 183)].**

| Scientific name | Common name | San Miguel | Santa Rosa | Santa Cruz |
|---|---|---|---|---|
| **Ferns & Fern Allies** | | | | |
| *Equisetaceae sp.* | scouring rush | | x | x |
| Unknown | unidentified ferns | x | x | x |
| **Conifers** | | | | |
| *Pinus muricata forma muricata* | Bishop pine | | x | x |
| *Pinus torreyana s. insularis* | Santa Rosa Island Torrey Pine | | x | |
| **Dicotyledonous Flowering Plants** | | | | |
| **Aizoaceae** | **Iceplant Family** | | | |
| *Carpobrotus sp.* | pigface | x | x | x |
| *Mesembryanthemum* | iceplant | x | x | x |
| **Anacardiaceae** | **Sumac Family** | | | |
| *Rhus integrifolia* | lemonade berry | x | x | x |
| *Rhus ovata* | sugar bush | | | x |
| *Toxicodendron diversilobum* | Pacific poison oak | x | x | x |
| **Apiaceae** | **Celery or Carrot Family** | | | |
| *Foeniculum vulgare* | sweet fennel | | x | x |
| *Daucus pusillus* | rattlesnake weed | x | x | x |
| **Asteraceae** | **Sunflower Family** | | | |
| *Achillea millefolium* | common yarrow | x | x | x |
| *Artemisia californica* | coastal sagebrush | x | x | x |
| *Aster sp.* | aster | | x | x |
| *Baccharis pilularis s. consanguinea* | coyote brush | x | x | x |
| *Baccharis salicifolia* | mulefat | x | x | x |
| *Cirsium sp.* | thistle | x | x | x |
| *Coreopsis gigantean* | giant coreopsis | x | x | x |
| *Eriophyllum confertiflorum* v. *confertiflorum* | golden yarrow | x | x | x |
| *Isocoma sp.* | coastal goldenbush | x | x | x |
| *Malacothrix sp.* | dandelions | x | x | x |
| **Brassicaceae** | **Mustard Family** | | | |
| *Brassica rapa* | field mustard | x | x | x |

| | | | | |
|---|---|---|---|---|
| **Cactaceae** | **Cactus Family** | | | |
| *Opuntia sp.* | prickly pear | x | x | x |
| **Caryophyllaceae** | **Pink Family** | | | |
| *Silene sp.* | pink | x | x | x |
| **Chenopodiaceae** | **Goosefoot Family** | | | |
| *Atriplex sp.* | saltbush | x | x | x |
| *Chenopodium* | goosefoot | x | x | x |
| **Convolvulaceae** | **Morning Glory Family** | | | |
| *Calystegia macrostegia* s. *macrostegia* | Northern island morning glory | x | x | x |
| **Crassulaceae** | **Stonecrop Family** | | | |
| *Dudleya sp.* | dudleya | x | x | x |
| **Fabaceae** | **Pea Family** | | | |
| *Astragalus sp.* | locoweed | x | x | x |
| *Lathyrus vestitus* v. *vestitus* | wild sweet pea | | x | x |
| *Lotus sp.* | deerweed, lotus | x | x | x |
| *Lupinus sp.* | lupine | x | x | x |
| *Trifolium sp.* | clover | x | x | x |
| *Vicia sp.* | vetch | x | x | x |
| **Fagaceae** | **Oak Family** | | | |
| *Quercus sp.* | oak | | | x |
| **Myrtaceae** | **Myrtle Family** | | | |
| *Eucalyptus sp.* | gum | | | x |
| **Nyctaginaceae** | **Four-o'-clock Family** | | | |
| *Abronia sp.* | sand-verbena | x | x | x |
| **Onagraceae** | **Evening Primrose Family** | | | |
| *Camissonia sp.* | primrose | x | x | x |
| *Clarkia sp.* | misc. flowering plants | x | x | x |
| *Epilobium sp.* | fuchsia | x | x | x |
| **Papaveraceae** | **Poppy Family** | | | |
| *Eschscholzia californica* | California poppy | x | x | x |
| Unknown | unidentified poppies | x | x | x |
| **Polygonaceae** | **Buckwheat Family** | | | |
| *Eriogonum sp.* | buckwheat | x | x | x |
| *Rumex sp.* | dock | x | x | x |
| **Portulacaceae** | **Purslane Family** | | | |
| *Calandrinia sp.* | misc. flowering plants | x | x | x |

| | | | | |
|---|---|---|---|---|
| *Claytonia sp.* | Miner's lettuce | x | x | x |
| **Primulaceae** | **Primrose Family** | | | |
| *Dodecatheon clevelandii* | shooting star | x | | |
| **Ranunculaceae** | **Buttercup Family** | | | |
| *Ranunculus californicus* | California buttercup | x | x | x |
| **Rhamnaceae** | **Buckthorn Family** | | | |
| *Ceanothus arboreus* | island ceanothus | | x | x |
| *Ceanothus megacarpus* | big-pod ceanothus | | x | x |
| *Rhamnus californica* | coffee berry | | | |
| *Rhamnus pirifolia* | island redberry | | | x |
| **Rosaceae** | **Rose Family** | | | |
| *Adenostoma fasciculatum* | chamise | | x | x |
| *Heteromeles arbutifolia* | toyon | x | x | x |
| *Prunus ilicifolia s. lyonii* | island cherry | | x | x |
| *Rosa californica* | California wild rose | | | x |
| **Salicaeae** | **Willow Family** | | | |
| *Populus sp.* | cottonwood | | x | x |
| *Salix sp.* | willow | x | x | x |
| **Scrophulariaceae** | **Figwort Family** | | | |
| *Castilleja sp.* | paintbrush | x | x | x |
| *Castilleja sp.* | clover | x | x | x |
| *Mimulus sp.* | monkeyflower | | x | x |
| **Solanaceae** | **Nightshade Family** | | | |
| *Lycium sp.* | boxthorn | x | x | x |
| **Verbenaceae** | **Vervain Family** | | | |
| *Verbena lasiostachys* | verbena | x | x | x |
| **Monocotyledonous Flowering Plants** | | | | |
| **Alliaceae** | **Onion Family** | | | |
| *Allium praecox* | early onion | x | x | x |
| *Dichelostemma capitatum* | blue dicks | x | x | x |
| **Cyperaceae** | **Sedge Family** | | | |
| *Carex sp.* | sedge | | x | x |
| **Iridaceae** | **Iris Family** | | | |
| *Sisyrinchium bellum* | blue-eyed grass | x | x | x |
| **Juncaceae** | **Rush Family** | | | |
| *Juncus sp.* | rush | x | x | x |
| **Liliaceae** | **Lily Family** | | | |

| | | | | |
|---|---|---|---|---|
| *Calochortus albus* | fairy lanterns | | x | x |
| *Zigadenus fremontii* | death-camas | x | x | x |
| **Poaceae** | **Grass Family** | | | |
| *Achnatherum diegonense* | San Diego needlegrass | x | x | x |
| *Avena sp.* | oats | x | x | x |
| *Bromus sp.* | brome | x | x | x |
| *Hordeum sp.* | barley | x | x | x |
| *Hordeum murinum* | foxtail | x | x | x |
| *Nassella pulchra* | purple needlegrass | x | x | x |
| *Polypogon monspeliensis* | rabbitsfoot grass | x | x | x |
| *Vulpia sp.* | fescue | x | x | x |
| **Typhaceae** | **Cattail Family** | | | |
| *Typha sp.* | cattail | | x | x |

**Figure S1.** Experimental chambers for in-situ foraging efficiency trials conducted on song sparrows from San Miguel and Santa Cruz Islands during the breeding season (Feb-June) in 2014. Trial cages (44.5 cm x 22.2 cm x 26.7 cm) were equipped with two perches at different heights and absorbent material to collect excess fecal matter and minimize stress. We provided a plastic, visual block on one of the four sides of each cage to protect subjects from potential predators and extreme weather conditions which may influence stress and willingness to forage (Clinchy et al., 2013; Lima and Dill, 1990). We secured a translucent cloth on three of the remaining four sides to allow for light to enter thereby stimulating activity. The plastic shield contained a hole fitted to the lens of high definition Sanyo Dual Camera Xacti WH1 camcorders (Sanyo Model VPC-WH1YL), which were mounted on the outer side of the shield on flexible tripods.

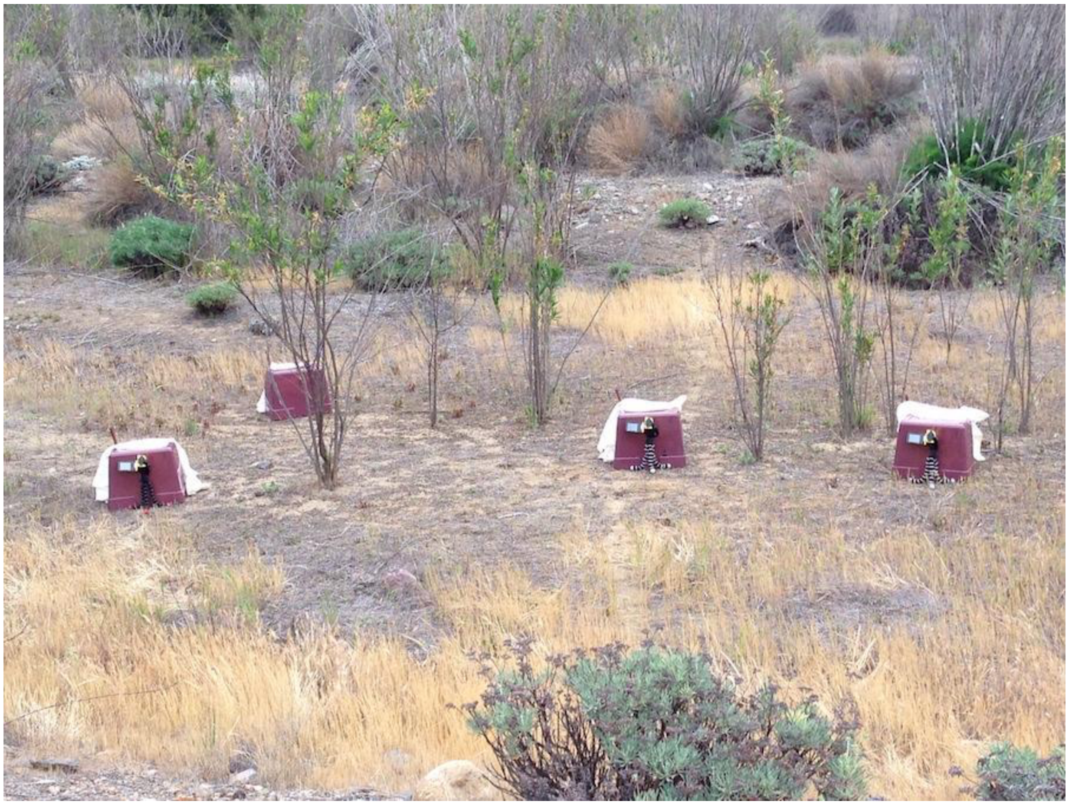

**Figure S2. Variable loadings of principal component analyses (PCA) of bill dimensions [depth (billd), length (billl), and width (billw)] for analyses of foraging efficiency.** PCA was conducted on bill dimensions in 33 individuals (San Miguel Island, *n* = 10; Santa Cruz Island, *n* = 23) to generate a composite score of overall bill morphology. Bill depth and width loaded equally (44%), and bill length did not strongly load on PC1 (12%).

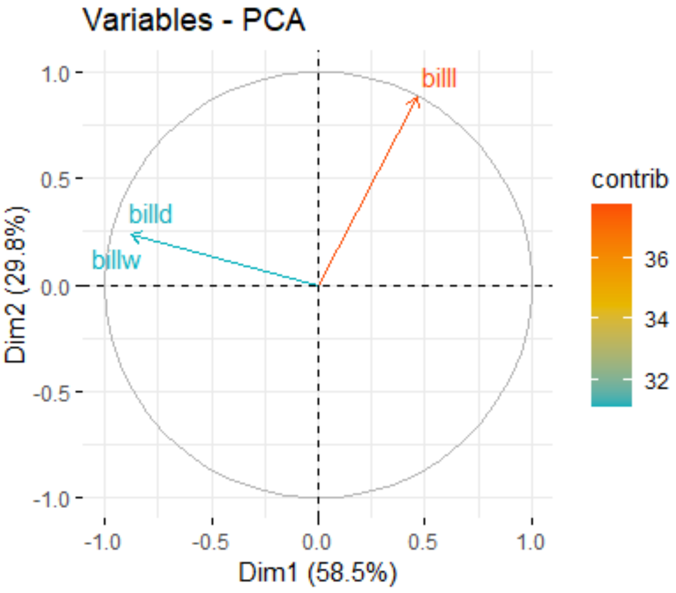

**Figure S3. Results from power analyses for bite force.** We performed power analyses to test if sample sizes were sufficient to detect biologically relevant differences between paired populations. We predicted biologically relevant differences in bite force would be 1.5 Newtons. We simulated data based on our means and variances measured in this study and then compared the model fit following Bolker et al. (2009).

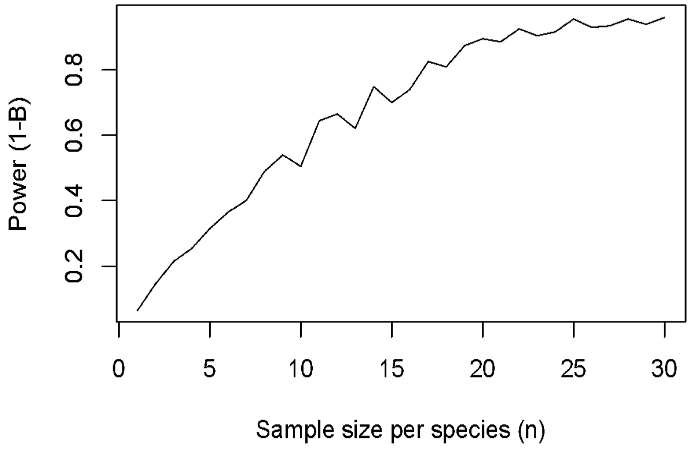

### REFERENCES

**Bolker, B. M., Brooks, M. E., Clark, C. J., Geange, S. W., Poulsen, J. R., Stevens, M. H. H. and White, J.-S. S.** (2009). Generalized linear mixed models: a practical guide for ecology and evolution. *Trends Ecol. Evol.* **24**, 127–35.

**Clinchy, M., Sheriff, M. J. and Zanette, L. Y.** (2013). Predator-induced stress and the ecology of fear. *Funct. Ecol.* **27**, 56–65.

**Lima, S. L. and Dill, L. M.** (1990). Behavioral decisions made under the risk of predation: a review and prospectus. *Can. J. Zool.* **68**, 619–640.
